# Faded visual afterimages reappear after TMS over early visual cortex

**DOI:** 10.1101/383893

**Authors:** Engelen T., Rademaker R.L., Sack A.T

**Affiliations:** Department of Cognitive Neuroscience, Maastricht University, Maastricht, The Netherlands; Département d’Études Cognitives, École Normale Supérieure – PSL university, Paris, France; Psychology department, University of California San Diego, La Jolla, California, USA; Donders Institute for Brain, Cognition and Behavior, Radboud University, Nijmegen, the Netherlands

## Abstract

In the complete absence of small transients in visual inputs (e.g. by experimentally stabilizing an image on the retina, or in everyday life during intent staring), information perceived by the eyes will fade from the perceptual experience. While the mechanisms of visual fading remain poorly understood, one possibility is that higher-level brain regions actively suppress the stable visual signals via targeted inhibitory feedback onto Early Visual Cortex (EVC). Here, we used positive afterimages and multisensory conflict to induce gestalt-like fading of participants’ own hands. In two separate experiments, participants rated the perceived quality of their hands both before and after Transcranial Magnetic Stimulation (TMS) was applied over EVC. In a first experiment, triple pulse TMS was able to make a faded hand appear less faded after the pulses were applied, compared to placebo pulses. A second experiment demonstrated that this was because triple pulse TMS inoculated the removed hand from fading over time. Interestingly, TMS similarly affected the left and right hand, despite being applied only over right EVC. Together, our results suggest that TMS can lift inhibitory processes in EVC and reverse the effects of visual fading. And it might do so by crossing transcollosal connections, or via multimodal integration sites in which both hands are represented.

## Introduction

If you’ve ever stared steadily enough at a paint chip on a wall, or a crack in the ceiling, you may have noticed the peripheral world slowly fading into oblivion. This visual fading, known as the Troxler effect (Troxler, 1804) is one of multiple examples of visual fading on record (Billock & Tsou, 2004; Ditchburn & Ginsborg, 1952; Riggs, Ratliff, Cornsweet, & Cornsweet, 1953). It is generally understood that small transients in the visual input, such as those induced via microsaccades, are critical to establish a visual percept from light entering the eyes, while images that are stable on the retina will result in the visual world fading away (Martinez-Conde, Macknik, & Hubel, 2004; Martinez-Conde, Otero-Millan, & MacKnik, 2013). While the phenomenon of visual fading has a long history in experimental psychology (Darwin, 1795), its neural substrate remains poorly understood. One potential mechanism is that stable visual signals are actively inhibited at the level of Early Visual Cortex (EVC). As a direct test of this hypothesis, here we used Transcranial Magnetic Stimulation (TMS) over EVC while a retinally stable afterimage faded from view. We demonstrated that visual fading is not inevitable, and can be partly reversed by EVC stimulation.

It is widely recognized that visual percepts are shaped by more than solely the input originating from the retina. This is demonstrated by illusions revealing a mismatch between what people report seeing, and the information entering their eyes. For example, during perceptual filling-in, information is perceived that is not directly sensed by the eyes (Komatsu, 2006; Shimojo, Kamitani, & Nishida, 2001). In contrast, perceptual fading is an example of an image sensed at the level of the retina, while that image is not being perceived (Billock & Tsou, 2004; Martinez-Conde et al., 2004, 2013). The subjective percept, and not retinal input, seems to determine responses at the earliest levels of visual processing. In perceptual filling-in, fMRI activity in primary visual area V1 tracked the subjectively perceived filled-in surface between two moving gratings (Meng, Remus, & Tong, 2005), and an illusory colored surface (Sasaki & Watanabe, 2004), rather than the objectively presented blank gap. Similarly, modulations of retinotopic activity in V1 have been demonstrated under conditions of constant retinal inputs: Retinotopic activity tracked the perceived size changes of retinally identical afterimages (Sperandio, Chouinard, & Goodale, 2012), and objects that appeared to occupy more space in the visual field (but had the same angular size) also activated a greater portion of V1 than objects that appeared to occupy less space (Murray, Boyaci, & Kersten, 2006). Finally, lower levels of visuo- cortical activity have been observed when a visual image (a monochromatic disk) faded into the background, compared to when it was consciously perceived (Mendola, Conner, Sharma, Bahekar, & Lemieux, 2006). Thus, conscious percepts during visual illusions are accompanied by changes at the earliest levels of sensory processing.

It is common for perceptual fading to occur in meaningful chunks, following Gestalt-like principles (Billock & Tsou, 2004). This implies the involvement of higher-level areas, able to process information at a more semantic level. Indeed, the neural correlates of the fading monochromatic disk extended beyond V1, and were also observed in higher-level areas including parietal cortex (Mendola, et al., 2006). Support for a causal link between parietal cortex and conscious visual percepts comes from a study showing that TMS stimulation over Inferior Parietal Sulcus (IPS) was able to elicit visual fading of a peripherally presented target. Presumably, TMS led to visual fading because it degraded the quality of attentional feedback signals to EVC (Kanai, Muggleton, & Walsh, 2008).

Given the interplay between both higher- and lower level areas of the brain, the question arises which neural mechanisms may underlie perceptual fading. Because changes in early sensory cortex generally track perception, here we hypothesize that dynamics at the level of EVC are responsible for visual fading. Fading might ensue when ongoing but stable sensory inputs are actively filtered out of the perceptual experience via inhibition. Because visual fading obeys Gestalt principles, targeted top-down inputs to sensory cortex might be a means to impose the early-level changes required for the perceptual experience of gestalt-like fading in afterimages.

To test this hypothesis, we utilized positive afterimages to induce perceptual fading. Positive afterimages can be elicited by discharging a brief flash of light after a prolonged period of dark adaptation. In the total darkness after the flash, a grey-scale visual afterimage develops of everything that was perceived during the flash. The experience can be described as a dim light turning on, with lights and darks having the correct sign (i.e. contrast is not inverted).

Positive afterimages are an elegant means to achieve full retinal stabilization of a scene and, importantly, they lend themselves well for inducing a conflict between vision and proprioception (Davies, 1973). In everyday life, visual and proprioceptive modalities will generally be in congruence with one another, i.e. when reaching for an object you will both see your hand reaching towards the object, as well as receive proprioceptive information on the trajectory of your hand. When viewing a positive afterimage, a situation can be created during which these two modalities convey incongruent messages to the brain. For example, when an afterimage is elicited while one is viewing one’s own hand, and subsequently that hand is moved out of the afterimage, the hand will appear to fade or crumble while the rest of the afterimage stays intact (Davies, 1973; Gregory, Wallace, & Campbell, 1959). In this case, proprioception has a catastrophic impact on visual perception when conflicting information is encoded from the two senses (i.e. proprioception indicates the hand is removed, while the retinal input indicates the hand is still in view). Note that this resolution to the sensory conflict is somewhat unique, as in most cases a conflict between vision and proprioception is resolved in favor of the visual sense (Hay, Pick, & Ikeda, 1965), presumably due to its high spatial acuity and reliable signals.

During perception in positive afterimages, there is a clear impact on visual processing that comes from the sense of proprioception. For example, when a person moves their hand nearer or further in a positive afterimage such movements can induce a perceived change in size: a hand will appear to shrink or grow, depending on the direction of the movement (i.e. towards or away from the eyes, respectively; Carey & Allan, 1996). Both fading and size scaling of a hand in an afterimage occur irrespective of whether the movement is active or passive, suggesting that afferent proprioceptive information is sufficient to influence the visual percept (Bross, 2000; Hogendoorn, Kammers, Carlson, & Verstraten, 2009).

The sensory conflict evoked when removing a hand from a positive afterimage, and the subsequent gestalt-like fading of the removed hand, could result from targeted feedback from multimodal areas onto EVC (Macaluso et al., 2000; Bolognini & Maravita, 2007). Specifically, fading of the hand could be due to the active inhibition of the bottom-up retinal signals associated with viewing that hand. While this type of fading is necessarily instantiated by signals originating from outside EVC, it might be sustained via continued feedback from higher-level areas, or via local mechanisms at the level of EVC. In either case, we hypothesize that active inhibition at the level of EVC is responsible for the perceptual experience of a hand fading in an afterimage. In fact, active inhibition might more broadly serve as the mechanism responsible for perceptual fading of stabilized images: In the absence of visual transients, information might be deemed irrelevant by the visual system, and hence systematically filtered out of the perceptual experience.

Here, we tested this hypothesis by combining fading in positive afterimages with online TMS over right EVC. If perceived gestalt-like fading of a hand is indeed the result of targeted inhibition at the level of EVC, disrupting this inhibition by means of TMS should weaken the extent to which the hand is perceived to fade (i.e. lead to a temporary ‘recovery’ of the faded hand). In two separate experiments, participants viewed afterimages of both their hands held out in front of them. In Experiment 1, one of the hands was removed, and participants rated the relative clarity of their two hands immediately before, and immediately after, a TMS-manipulation over right EVC. Manipulations consisted of a single TMS pulse, a triple TMS pulse, or a single placebo TMS pulse. In Experiment 2 only triple pulses (placebo and real TMS) were administered, and participants rated both hands individually to disentangle the effects of TMS on hands ipsilateral and contralateral to the targeted right hemisphere. In both experiments we demonstrated that the gestalt-like fading of the removed hand was systematically reduced after administering TMS over EVC. This implies that TMS can provide temporary relief from the factors responsible for perceptual fading, and suggests that active inhibitory mechanisms impacting visual processing at the level of EVC are likely at play during perceptual fading.

## Methods

### Participants

Participants in Experiment 1 were ten healthy volunteers recruited from Maastricht University with a mean age of 25.8 years (SE = 3.3, 8 females). Ten more participants took part in Experiment 2, of whom two had previously participated in Experiment 1 (mean age 25.5 years, SD = 2.8; 6 females). All participants were right handed, had normal or corrected-to-normal vision, and were naïve to the purpose of the experiment (except for author RR in Experiment 2). Before the start of the experimental proceedings, participants provided written informed consent and were screened for TMS safety based on published safety guidelines (Rossi, Hallett, Rossini, & Pascual-Leone, 2012). In total, we excluded nine people from participation on the grounds that they were unable to perceive phosphenes (N = 2), unable to perceive phosphenes at the desired visual field location (N = 1), unable to experience an afterimage long enough to give the ratings within the time the image was perceived (N = 2), or unable to experience fading in the afterimage in response to bodily movement (see also procedures, N = 4). This study was approved by the standing ethical committee of the Psychology and Neuroscience department at Maastricht University. Except for one of the authors, participants received monetary reimbursement for their time.

### Materials

The experiments took place in a completely darkened room. Because some scattered light could not be avoided, a large cardboard box (65 x 74 x 95 cm) placed atop a table was used to enclose participant’s field of view and shield them from stray light, thus ensuring absolute darkness (Figure 1). Head and TMS coil stability were ensured by a forehead- and chinrest, together with a coil-holder firmly fixing the TMS coil over the participant’s skull. To allow stable fixation in the dark, dim red LED light was shone through a small hole in the box, creating a fixation of < 0.25° of visual angle at a viewing distance of 55cm. The wavelength of red light (620-750 nm) falls outside of the response range of retinal rods (Wald, 1955), and as such did not interfere with dark adaptation. To ensure stable arm and hand positions from one trial to the next, a horizontal bar was placed at ~22cm above the tabletop, at 42 cm from the front of the box. This distance required participants to raise their elbows up from the tabletop to place their hands behind the bar. Participant’s arms were obscured from view by means of a black plastic arm-occluder attached to the bar, and the lateral hand distance was fixed by two markers behind the bar that were readily discernable by touch. To elicit positive afterimages, a Vivitar 285HW Zoom Thyristor flashgun was placed in the box pointing at the white box-ceiling above the participant.

**Figure 1.**
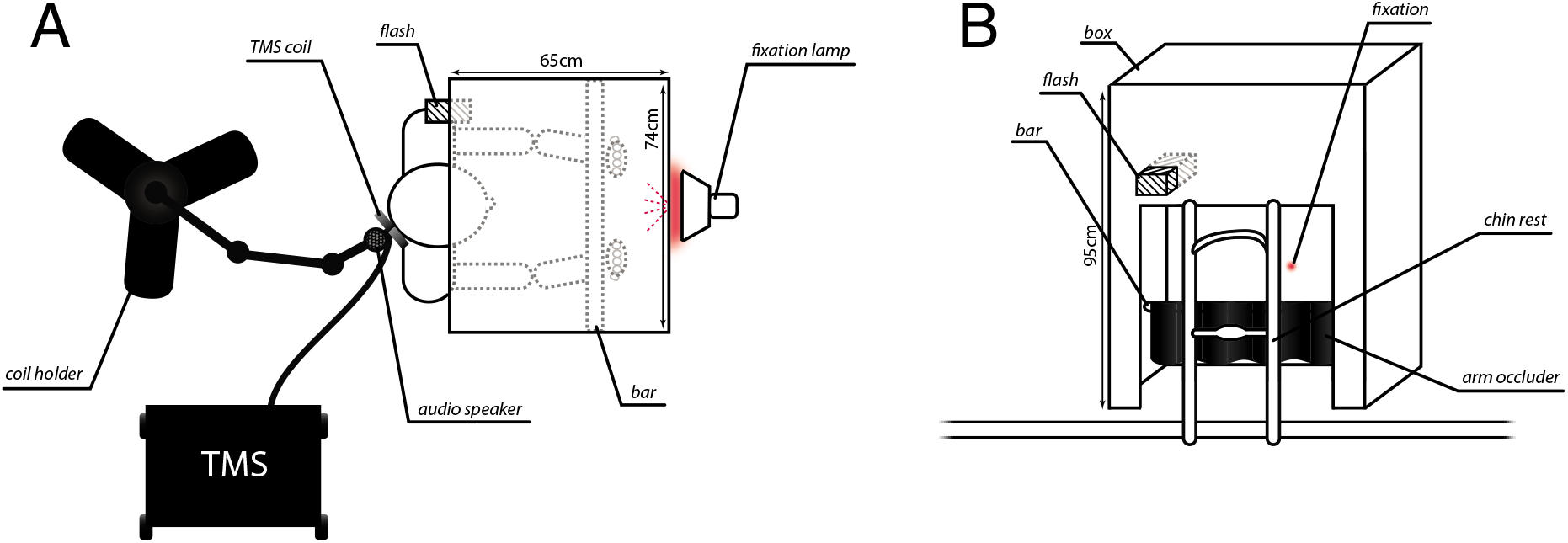
Experimental setup. (A) Top view of cardboard box with participant seated in the experimental setup. Stimulator and coil-holder are also depicted. (B) Front view of cardboard box atop of the table, depicting also the chinrest and fixation. Images are scaled to true size.

Biphasic TMS stimulation was applied with a MagPro R30 stimulator (Medtronic Functional Diagnostics A/S, Skovlunde, Denmark; maximum stimulator output 1.9T) and a figure-of-eight coil (MCB70). The coil was placed with the handle oriented laterally to the right and held throughout the experimental procedures by a custom-made coil-holder, ensuring stable placement over the skull. Auditory placebo pulses were delivered over an audio speaker that was attached to the coil-holder, adjacent to the TMS coil (Figure 1A). TMS sounds, as emitted by the TMS coil, were recorded with an iRiver recording device and edited offline to filter out background noise with the Audacity software package. Volume (in dB) was measured with a decibel meter for both placebo (generated by the speaker) and real (generated by the TMS coil) pulses across a range of intensities, and the volume of the placebo pulses was equated to match the volume of the real pulses. To control trial timing in the dark, auditory beeps indicated the start of every new trial at 40-second intervals. Beeps were played from a computer, and were generated using the Presentation software package (Experiment 1) and Matlab with the Psychophysics toolbox (Experiment 2; Brainard, 1997).

## Procedure

Each of the two experiments consisted two separate sessions: A preliminary and an experimental session. The purpose of the preliminary session was (1) to check if participants could perceive phosphenes, (2) to familiarize participants with afterimages and experimental procedures, (3) to practice the rating scale, and (4) to screen participants, and see whether movement of a hand resulted in perceived fading of that hand from the afterimage. This screening procedure was adapted from previous work (Carlson, Alvarez, Wu, & Verstraten, 2010; Rademaker, Wu, Bloem, & Sack, 2014) and required because both Experiments 1 and 2 critically depended on participant’s ability to perceive fading in response to bodily movements.

During the preliminary session participants were seated with their head in the chinrest, and instructed to fixate throughout. This session consisted of 15 practice trials performed after 10 minutes of dark adaptation: The first 3 trials were to familiarize participants with afterimages, without performing a task. During the following 6 trials, participants placed their hands behind the horizontal bar with their palms facing towards them. After the flash, the left or right hand was removed (or both were kept stationary in experiment 2), and participants described their percept. During the final 6 trials participants practiced quantifying their subjective fading experiences, using the rating scale. For Experiment 1 we used a rating scale that relied on a relative judgment of the two hands on a 11- point scale (adapted from Hogendoorn, Kammers, Carlson, & Verstraten, 2009), and for Experiment 2 we used a variant of this scale that allowed the hands to be judged independently on a 7-point scale (Figure 2C and Figure 2D respectively). The preliminary session took approximately 1 hour to complete. Some participants were allowed extra practice trials in case of difficulties mastering the task and/ or rating scale.

**Figure 2.**
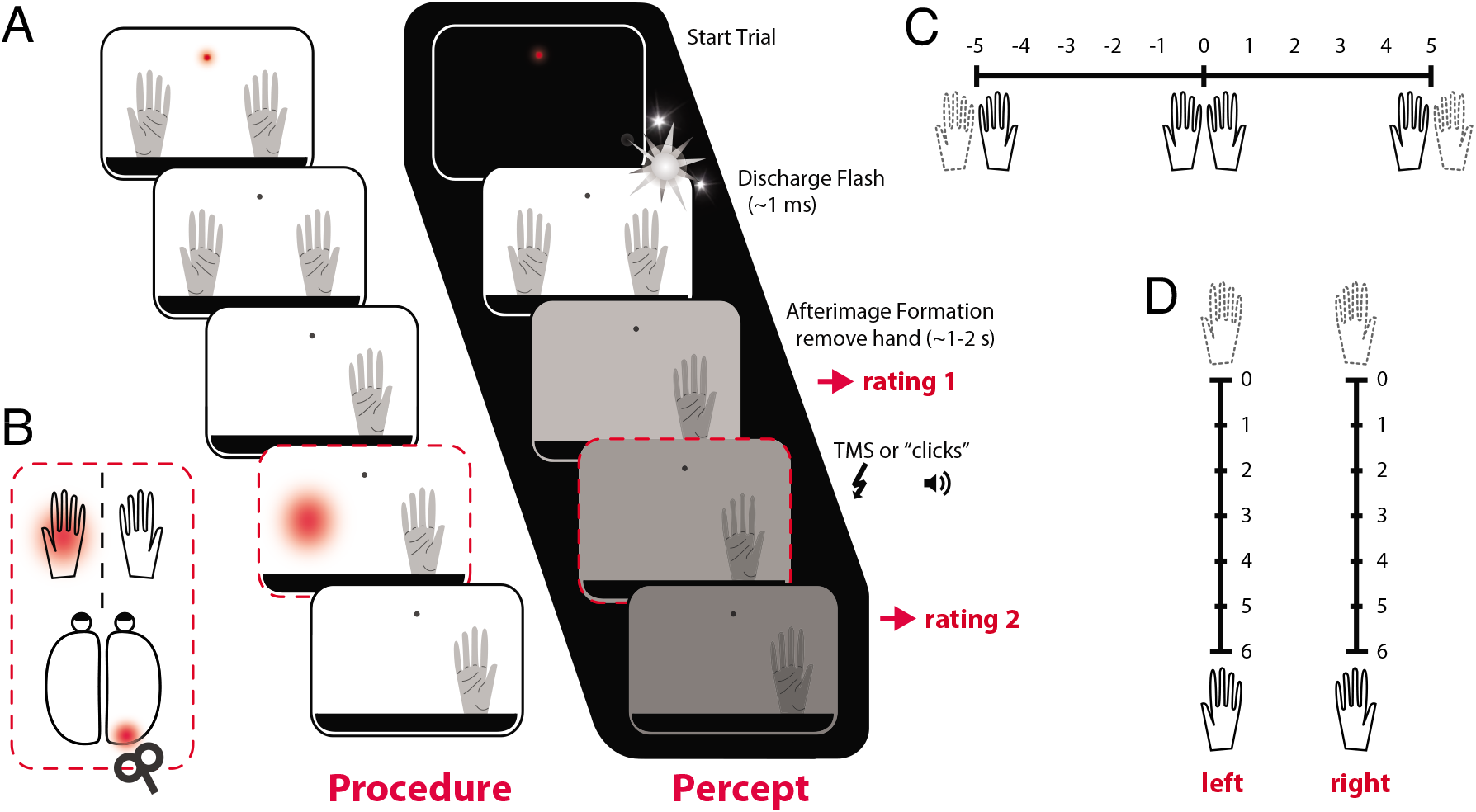
Trial sequence, stimulation, and rating scales. (A) At the start of each trial, participants put their hands up behind the black bar and started fixation aided by a small red light. When participants indicated that they were ready to start the trial, a brief flash was emitted, and once the afterimage had developed participants removed one of their hands (or kept both stationary). After briefly observing their hands, participants gave a rating, which triggered placebo (auditory clicks) or TMS pulses. After the pulses, participants again briefly observed their hands and provided a second rating. The next trial started 40 seconds after the start of the previous trial, and in between trials participants could rest their arms, and move their eyes around freely. (B) TMS was applied over the right part of Early Visual Cortex corresponding to the left part of visual space. To stimulate the part of the visual field containing the left hand, prior to the start of the experiment phosphenes were elicited and the coil moved around until the left hand and phosphenes overlapped. During the experiment itself, stimulation was applied at 80% of phosphene threshold to ensure participants would not perceive any phosphenes. (C) Rating scale Experiment 1. A rating of zero would indicate that both hands were perceived with equal strength. A rating of -5 indicated that the left hand had faded entirely, while the right hand was perceived as veridical. A rating of 5 meant the opposite, with the left hand being perceived as veridical, and the right hand having faded completely. (D) Rating scale Experiment 2. Half of the participants would always rate first the left, then the right hand, while the other half of participants always rated first the right, then the left hand.

An experimental session started by positioning the coil over the right posterior part of the skull, and identifying the part of early visual cortex corresponding to the visual field position of the left hand by means of phosphene localization (Figure 2B). Once properly localized, the coil was fixed over the skull with a coil- holder, and participant’s phosphene thresholds were determined in an already dimmed room, using a brief staircase procedure. Subsequent stimulation was applied at 80% of phosphene threshold (mean = 31.7% of stimulator output with SD = 4.8% in experiment 1, and mean = 39.2% of stimulator output with SD = 10.9% in experiment 2). After ten minutes of dark adaptation, participants completed 6 blocks of 15 trials per block. A trial (Figure 2A) started by participants placing their hands behind the bar, and after the flash was emitted and an afterimage had formed, participants removed one of their hands (or kept both up) and after a brief period of observation gave a rating. This first rating triggered the TMS or placebo pulses (Figure 2A), after which participants again observed and rated their hands. Participants fixated throughout each trial. To avoid drift of the afterimage relative to fixation, the red light was always switched off simultaneously with the flash, after which participants continued to fixate the unlit fixation spot. Participants could rest their arms on the tabletop in between trials.

The six experimental blocks of Experiment 1 consisted of two conditions during which a hand was removed (either the left or right hand), and three stimulation conditions (either a single placebo sound pulse, a single pulse TMS, or triple pulse TMS at 10Hz). Because TMS targeted the left part of visual space overlapping with the position of the left hand, this design allowed us to evaluate the effect of TMS on fading when the left hand was removed. Moreover, this could be compared to how TMS affected fading of the right hand, and to fading during placebo instead of real TMS. Experiment 2 consisted of three conditions involving the hands (left hand was removed, right hand was removed, or both hands remained stationary) and two stimulation conditions (triple placebo sound pulses, or real triple pulse TMS at 10Hz). The stationary condition allowed us to see how TMS affected the percept of a hand when there was only gradual fading of the entire afterimage occurring naturally over time. The order of blocks was counterbalanced between participants. An experimental session lasted between 1.5 and 2 hours.

## Results

### Experiment 1

Participants gave two ratings on a relative scale (Figure 2C) – one right before, and one right after the pulse. If the left hand were less visible than the right, participants gave a negative rating. Conversely, positive ratings would be given if the right hand were faded relative to the left. If both hands were equally visible, the rating would be “0”. Note that to equate the effects of removing the left and right hands (negative- and positive-going, respectively), we inverted the sign of the ratings given when the left hand was removed *before statistical testing*. Thus, one can consider the negative-going average ratings displayed in the top panel of Figure 3A as positive (and consequently, the positive-going light gray bars in Figure 3B as negative), for the purposes of interpreting all subsequent analyses reported here.

**Figure 3.**
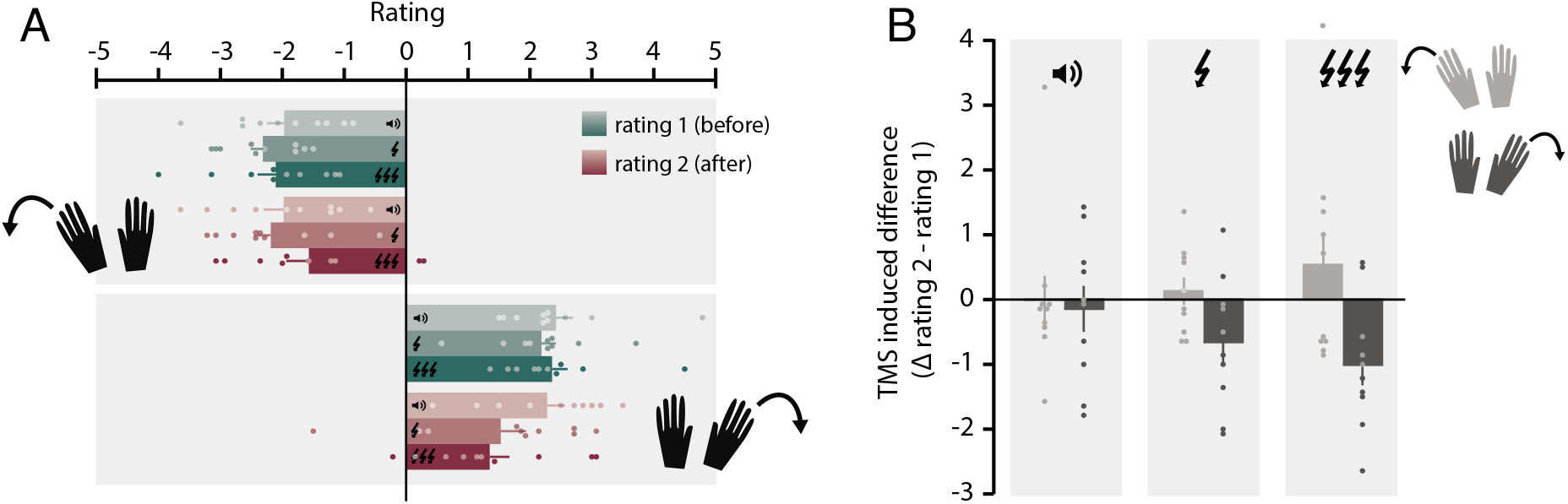
The effects of placebo and real TMS on fading of the left and right hands in Experiment 1. (A) Participants rate the relative visibility of their hand after removing one hand from view both before and after a pulse (placebo, single pulse, or triple pulse TMS). Before the pulse (green bars) there were no differences between pulse conditions. After the pulse there was a difference between the pulse conditions, with relative ratings closer to zero after triple pulse TMS compared to a placebo sound pulse. This indicates that after triple pulse TMS the two hands were rated as more similar in their degree of visibility. (B) Data in (A) are replotted to more clearly show the impact of the pulse by contrasting the ratings before and after the pulse. A difference of zero indicates no change in relative ratings before and after the pulse. For trials on which the left hand was removed (light grey) a positive score indicates that the pulse made the two hands appear more similar in terms of their visibility. Similarly, for trials on which the right hand was removed (dark grey) a negative score indicates that the pulse made the two hands appear more similar in terms of their visibility. Note that, to perform statistical testing, the scores given after left hand removal were inverted to equate the directionality of the fading effect.

To verify that participants experienced gestalt-like fading of the hand removed from the afterimage before the pulse (Figure 3A; green bars), we performed a 2 (hand) x 3 (TMS condition) repeated-measures ANOVA against zero. Removing a hand from the afterimage resulted in perceived fading of that hand (F(1,9) = 93.150, *p* < .001, η^2^_p_ = 0.912), with average ratings between 1.97 and 2.42 across conditions. Moreover, equal amounts of fading were experienced before the pulse, with no differences between the left and right hands (F(_1_,_9_) = 1.390, *p* = 0.269, η^2^p = 0.134), nor any differences between TMS conditions (F(2,18) = 0.118, *p* = 0.889, η^2^p = 0.013). Indeed, no differences were expected at this baseline rating, prior to the application of TMS.

Next, we investigated the impact of the pulses on perceived fading (compare green and red bars in Figure 3A). A 2 (hand) x 3 (TMS condition) x 2 (before and after rating) repeated measures ANOVA showed a significant interaction between TMS condition and before/after rating (F(2,18) = 3.940, *p* = 0.038, η^2^p = 0.304). We followed up on this interaction by first collapsing the data across the hand condition, as relative ratings were no different for the left and right hand (F(1,9) < 0.001, *p* = 0.997, η^2^p < 0.001). Second, to compare the before and after ratings, we performed 3 post-hoc t-tests, each within a given TMS condition. Perceived fading did not change between the first (before) and second (after) rating in the placebo condition (mean before = 2.20 (SE = 0.26), mean after = 2.13 (SE = 0.28); t(10) = 0.214, *p* = 0.835), nor was there a significant change with single pulse TMS (mean before = 2.25 (SE = 0.19), mean after = 1.86 (SE = 0.37); t(10) = 1.786, *p* = 0.108). When triple pulse TMS was applied, ratings after the pulses were marginally closer to zero than before TMS (mean before = 2.23 (SE = 0.26), mean after = 1.46 (SE = 0.28); t(10) = 2.197, *p* = 0.056). This indicates that the relative visibility of the two hands became more similar after triple pulse TMS, irrespective of whether the left or right hand was removed. Third, to compare TMS conditions we performed two repeated-measures ANOVA’s, one within the before, and one within the after ratings. As already reported above, there were no differences between TMS conditions at the first (before) rating (F(2,18) = 0.118, *p* = 0.889, η^2^p = 0.013). However, at the second (after) rating there was a significant main effect of TMS condition (F(_218_) = 6.221, *p* = 0.009, η^2^p = 0.409), driven by the difference between placebo and triple pulse TMS *(t(9)* = 3.02, *p* = 0.043, Bonferroni corrected).

Figure 3B replots the data in Figure 3A by showing the difference between the before and after ratings. Here, the further the difference score is away from zero, the more similar the hands became after the pulse (in positive and negative directions for removing the left and right hands, respectively). An analysis of the difference scores in a 2 (hand) x 3 (TMS condition) repeated-measures ANOVA revealed a significant main effect of pulse condition (F(2,18) = 3.960, *p* = 0.038, η^2^p = 0.306), but no effect of hand (F(1,9) = 1.338, *p* = 0.277, η^2^p = .129), and no interaction (F(2,18) = 0.827, *p* = 0.453, η^2^p = 0.084). The main effect of TMS condition was driven by a trending difference between placebo and triple pulse TMS *(t(9)* = 2.816, *p* = 0.06, Bonferroni corrected), indicating the two hands were perceived as more similar after a triple TMS pulse, compared to placebo. Together, these results suggest that triple-pulse TMS significantly reduced the amount of fading participants experienced after removing their hand from an afterimage, irrespective of the hand (the left or right) that was removed.

### Experiment 2

The relative rating scale employed in Experiment 1 comes with two important limitations. First, under the assumption that TMS over right EVC exclusively targets the location of visual space encompassing the left hand, the relative rating should reflect perceived changes at this location. Thus, the TMS induced reduction in relative rating when the left hand was removed implies that TMS *reinstated* the visual representation of the left hand. Following the same logic, the TMS induced reduction in the relative rating when the right hand was removed implies that TMS *reduced* the quality of the percept of the stationary left hand, making it more similar in quality to the faded right hand. However, we cannot exclude alternative explanations for the results in Experiment 1. For example, if TMS over right EVC were able to also affect perception of the right hand, relative ratings would not dissociate between what is happening at the two hands. To illustrate this point, consider a trial during which the left hand was removed: If the relative rating was closer to zero after applying TMS, this could mean that the perceived quality of the left hand was improved, or alternatively, that the quality of the right hand was reduced. Both would make the hands appear qualitatively more similar. A second limitation of the relative rating scale is that it’s not sensitive to detecting overall fading of hands in the afterimage: Two hands that were both very crisp would receive a rating of zero, while two hands that had nearly faded entirely, but to an equal extent, would also get a rating of zero.

To address these limitations, Experiment 2 required participants to rate the left and right hand independently, and on an absolute scale from 0-6 (where 0 indicated complete fading of the hand, and 6 the crispest possible afterimage of the hand). This allowed us to directly observe the effect of TMS on each individual hand, and also to track overall fading levels of the hands in the afterimage. Furthermore, Experiment 2 included a condition in which neither hand was removed, allowing us to observe potential effects of TMS on an afterimage in the absence of gestalt-like fading. Only triple placebo sound pulses and real triple pulse TMS were used in Experiment 2.

We first analyzed the data as we did in Experiment 1 to verify that our effects replicated. To this end, a relative rating was calculated by subtracting the ratings of the individual hands (left minus right; Figure 4). As before, we inverted the sign of the relative ratings for the left hand removal condition before statistical testing to equate the directionality of the fading effect between the two hands. We confirmed that removing both the left (F(1,9) = 13.740, *p* = 0.005, η^2^p = 0.604) and right (F(1,9) = 20.896, *p* = 0.001, η^2^p = 0.699) hand from the afterimage resulted in gestalt-like fading before the pulses were applied (Figure 4a, green bars; repeated-measures ANOVA’s against zero), and the extent of this fading was comparable between removing the left and right hand (F(1,9) = 0.510, *p* = 0.493, η^2^_p_ = 0.054). When both hands were kept up, the right hand was relatively less visible than the left before the pulse (*F*(1,9) = 7.781, *p* = 0.021, η^2^p = 0.464), as indicated by the positive-going relative difference. However, the relative ratings were only slightly above zero (0.17 and 0.22 for placebo and real TMS, respectively) and these results did not hold up in direct t-tests against zero (t(9) = 1.98, *p* = 0.079 and t(9) = 1.75, *p* = 0.11 for placebo and real TMS, respectively).

**Figure 4.**
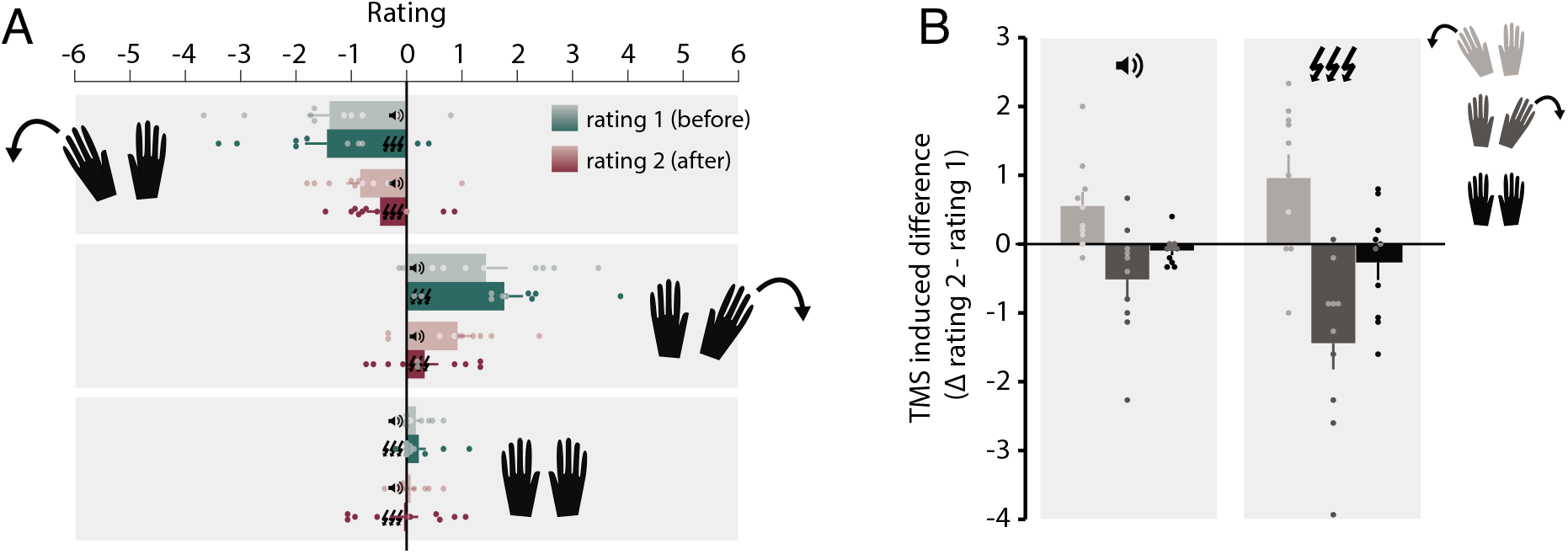
The effects of placebo and real TMS on fading of the left and right hand in Experiment 2. (A) Relative ratings of fading calculated from the absolute ratings participants had provided for individual hands. Note that the range of possible ratings in Experiment 2 (from -6 to 6) is slightly different from that in Experiment 1 (from -5 to 5). To ensure that both Experiments had an uneven number of rating options (with zero included), an 11-point scale was used in Experiment 1, and a 7-point absolute scale (i.e. 0-6, resulting in a 13-point relative scale) was used in Experiment 2. The effect of TMS pulses on fading was evaluated by calculating the difference between ratings (after minus before), as is shown in (B). A positive deflection along the y-axis indicates that ratings became more positive after TMS, as was the case when the left hand was removed (i.e. negative relative ratings before TMS would tend towards zero after TMS). Conversely, when the right hand was removed, a negative deflection along the y-axis indicates that ratings became more positive after TMS. Thus, the two hands were perceived as more similar after real TMS, irrespective of the hand that was removed from the afterimage.

We again assessed the differences between before and after ratings (Figure 4B). A 3 (hand) x 2 (TMS condition) repeated-measures ANOVA showed a main effect of hand condition (F(2,18) = 5.791, *p* = 0.011, η^2^p = 0.392), driven by a marginal difference between the both hands up and remove right conditions (t(_9_) = -2.845, *p* = 0.058), but not by any differences between the remove left and remove right conditions (t(_9_) = 1.383, *p* = 0.600). Importantly, ratings of the two hands became more similar after real TMS pulses than after placebo pulses (F(1,9) = 8.077, *p* = 0.019, η^2^p = 0.473), as indexed by a larger deviation away from zero on the y-axis for real TMS, which indicates that the two hands went from relatively different at rating 1 (before TMS), to relatively similar at rating 2 (after TMS). There was no significant interaction (F(2,18) = 3.164, *p* = 0.066, η^2^p = 0.260). These results replicate the findings of Experiment 1, insofar that real TMS caused perceived relative differences between a removed and a stationary hand to decrease compared to placebo.

Next, we analyzed the data from Experiment 2 by capitalizing on the absolute rating scale. Beyond what can be gleaned from relative difference scores, absolute ratings allowed us to assess fading at each hand individually. We first calculated the difference between the before- and after-ratings (Figure 5). A difference score of zero indicated no change in the perceived quality of the hand, while negative going scores indicated an increase in the amount of fading at the second (after) compared to the first (before) rating. Note that in Figure 5 all scores are negative going, indicating overall fading over time between the first and second rating.

**Figure 5.**
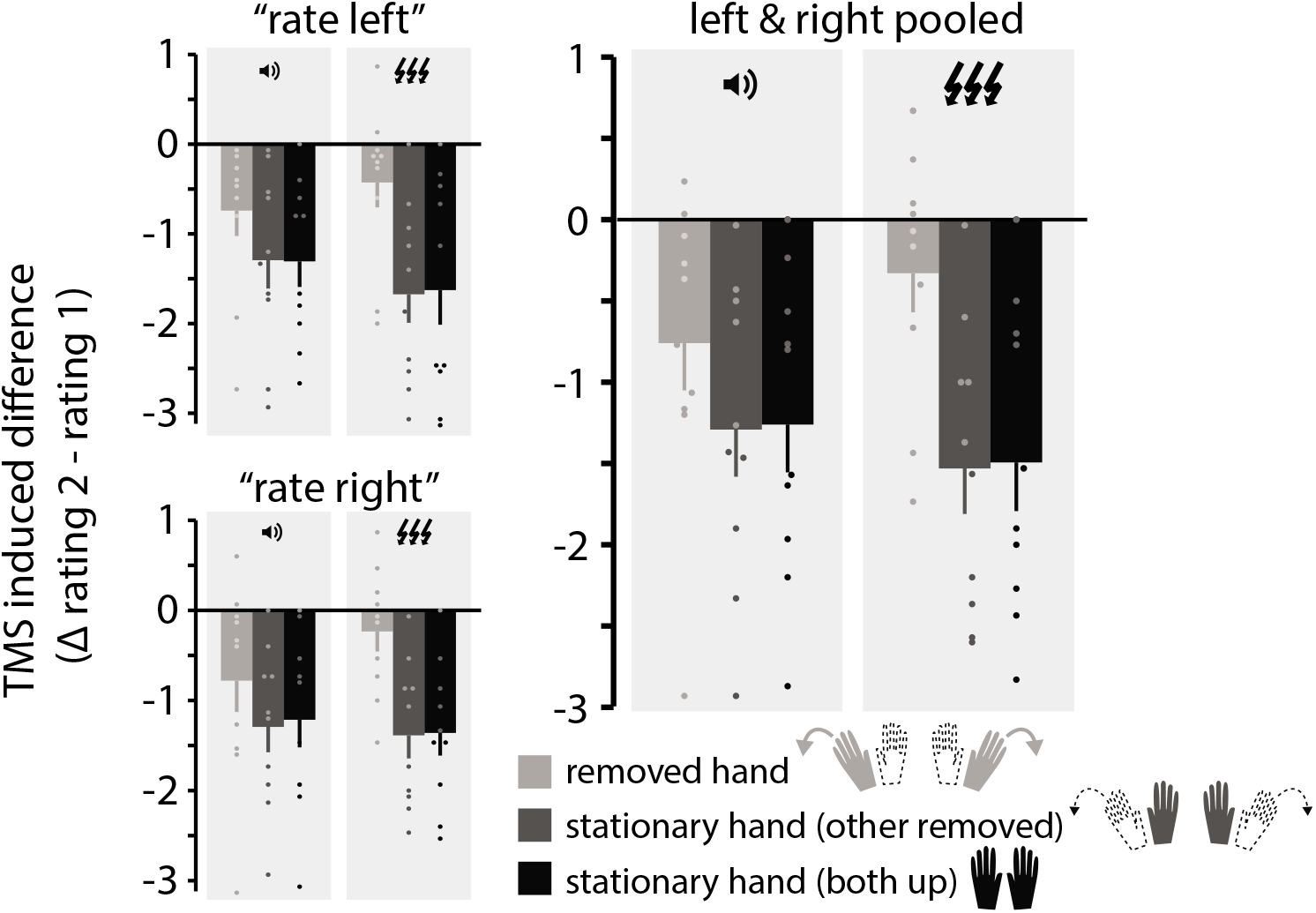
The effects of placebo and real TMS on fading of the left and right hands in Experiment 2. (A) Difference scores (rating after - rating before) are plotted against what occurred at the rated hand (i.e. 3 hand conditions). Hand condition was organized in three event types (1) the rated hand was removed, (2) the rated hand was stationary and the only hand kept up, and (3) the rated hand was stationary while both hands were kept up. More negative going scores indicate that the rated hand became less visible over time (i.e. between the first and second rating). (**B**) Replots the data in (A) pooled across the left and right hand.

For trials on which one of the hands was removed, we organized these difference scores according to the (in-) action at the rated hand. Specifically, we grouped difference scores by the rated hand being *removed* (i.e. right hand ratings when the right hand was removed, and left hand ratings when the left hand was removed) or *stationary* (i.e. right hand ratings when the left hand was removed, and left hand ratings when the right hand was removed). On trials where both hands were kept up, ratings of the left and right hand both refer to a situation in which a *stationary* hand was rated.

The results of a 3 (hand condition) x 2 (hand rated) x 2 (TMS condition) repeated-measures ANOVA showed no main effect of the hand being rated (*F*(_19_) = 2.002, *p* = 0.191, η^2^p = 0.182; Figure 5a), suggesting that any changes in response to the experimental manipulation occurred similarly in both the left and the right hand. Importantly, there was a significant interaction between hand condition and TMS condition (F(1,9) = 11.265, *p* = 0.001, η^2^p = 0.556). To explore this interaction, we first collapsed the data across the hand being rated (left or right). Next, we wanted to assess fading at the removed and stationary hands and, critically, how TMS influenced fading in the different hand removal conditions.

To assess fading when a hand was either removed or stationary, we ran two 3way repeated measures ANOVA, one for each TMS condition. Both ANOVAs revealed differences in fading between the three hand conditions (placebo: F(2,18) = 4.766, *p* = 0.022, η^2^p = 0.346; and real TMS: F(2,18) = 12.173, *p* < 0.001, η^2^p = 0.575). However, for placebo pulses none of the post-hoc pairwise comparisons remained significant (all *p* > 0.12), indicating that the differences in fading were not robust between the three hand conditions. Note also that in the removed hand condition there exists a ceiling effect: A removed and already faded hand has little room to fade further. Thus, the amount of fading that naturally occurs over time for the stationary hands is likely larger than the amount of fading still possible for the removed hand, also in the placebo condition. In contrast, post-hoc tests for real TMS showed significant differences between ratings of the removed hand and the stationary hand, both when the stationary hand was the only hand up (*t*(9) = 3.566, *p* = 0.018, Bonferroni corrected), as well as when both hands were up (*t*(9) = 3.566, *p* = 0.018, Bonferroni corrected).

Next, we aimed to assess how TMS condition influenced fading within the separate hand conditions, which was tested with three 2-way ANOVAs, one for each hand condition. When the hand was removed, placebo and real TMS had a different impact on perceived fading (F(1,9) = 5.417, *p* = 0.045, η^2^p = 0.376), with smaller scores for real (mean = -0.33 (SE = 0.24)) compared to placebo pulses (mean = -0.76 (SE = 0.29)). Because negative scores indicate that the hand faded from the first (before) to the second (after) rating, the real TMS score being more proximal to zero, compared to placebo pulses, suggests that real TMS inoculated the removed hand from fading.

When the hand was stationary, real TMS caused marginally more fading compared to placebo pulses, both when the stationary hand was the only hand up (F(1,9) = 4.606, *p* = 0.060, η^2^p = 0.339), and when both hands were up (F(1,9) = 4.046, *p* = 0.075, η^2^_p_ = 0.310). Scores were more strongly negative going with real TMS (mean = -1.53 (SE = 0.28) and mean = -1.49 (SE = 0.30) for one and both hands up, respectively) than with placebo pulses (mean = -1.29 (SE = 0.29), mean = -1.26 (SE = 0.29) for one and both hands up, respectively).

## Discussion

Activity in EVC correlates with subjective percepts, even when a percept is illusory and not a direct reflection of retinal inputs (Sasaki & Watanabe, 2004; Meng, Remus, & Tong, 2005; Murray, Boyaci, & Kersten, 2006; Sperandio, Chouinard, & Goodale, 2012). Such findings imply that visual fading occurs when bottom-up sensory information is muted, either via local inhibition or by targeted top-down inhibition. The role of top-down influences is supported by the fact that fading occurs in gestalt-like chunks (Billock & Tsou, 2004), hinting at the involvement of cortical areas with more complex response profiles. The experiments presented here set out to explore the mechanisms behind visual fading, and the hypothesized role of active inhibition at the earliest cortical stages of visual processing. Using gestalt-like fading of a hand removed from a retinally stabilized afterimage, we showed that online stimulation of EVC partially reversed visual fading. Thus, TMS stimulation over EVC could release the inhibition that exists otherwise in this region during visual fading.

In Experiment 1, participants removed either their left or right hand from a positive afterimage, after which the removed hand faded from view. Subsequently, participants rated the relative intensity of their two hands twice – directly before, and directly after single, triple, or placebo TMS. TMS was applied over the right EVC location corresponding to the visual field location occupied by the left hand. After triple pulse TMS, the two hands were rated as more similar, compared to after placebo or single pulse TMS. Curiously, the effect of triple pulse TMS over right EVC was not specific to the condition in which the left hand was removed (and faded) – the two hands were also rated as more similar after removal (and fading) of the right hand, positioned contralateral to the part of visual space targeted by TMS. Thus, triple pulse TMS impacted perception both when a hand contralateral to the targeted hemisphere was removed, and when a hand ipsilateral to the targeted hemisphere was removed. Notably, the rating scale used in Experiment 1 was based on the *relative* clarity of the two hands in the afterimage, obscuring the possible reason underlying this bilaterality. Specifically, when the right hand was removed, and the two hands were rated as more similar after triple pulse TMS, this could be because (1) an increase in the perceptual quality of the removed right hand (an effect ipsilateral to the TMS pulses), (2) a reduction of the perceptual quality of the stationary left hand (an effect contralateral to the TMS pulses), or (3) a combination of both.

In Experiment 2, participants removed their left or right hand from the afterimage, or kept both hands up. Participants then rated the *absolute* intensity of their two hands separately, both before and after three placebo or real TMS pulses. First, in an analysis aimed at replicating the relative ratings provided in Experiment 1, the application of triple pulse TMS resulted in ratings of the hands becoming more similar (compared to placebo pulses), indeed replicating our findings from

Experiment 1. Second, the absolute ratings at the individual hands further allowed us to evaluate the source of this effect. We found that triple pulse TMS resulted in a relative strengthening of the percept of both the left, as well as the right hand. These findings imply that TMS over right EVC can help reverse the effects of visual fading, and that the impact of the pulses is not hemifield specific.

Why would triple pulse TMS over right EVC, aimed at the visuo-spatial location of the left hand, reduce fading at both the left and right hands? First, TMS stimulation could have spread to the contralateral hemisphere through monosynaptic transcollosal connections (Berlucchi, & Rizzolatti, 1968). Indeed, previous work combining EEG with TMS over occipital cortex has demonstrated that a single TMS pulse to one hemisphere can spread to the contralateral hemisphere within 28ms after stimulation (Ilmoniemi et al., 1997). Given that we applied multiple pulses, and studied an effect that evolves on a timescale of seconds, rather than milliseconds, it is possible that the observed bilateral effects were due to inter-hemispheric spreading of TMS pulses.

Second, the effect of TMS over EVC might propagate to upstream areas with large visual receptive fields that can encompass the spatial location of both hands. If such areas in turn provide feedback to EVC, their ability to send inhibitory signals – to either of the removed hands – could be disrupted. In our paradigm, visual fading of a hand arises from a multimodal conflict between vision and proprioception. A putative mechanism for modulation of EVC activity through other modalities is via back-projections originating in multimodal integration areas, such as posterior parietal cortex (Macaluso et al., 2000; Bolognini & Maravita, 2007).

The posterior parietal cortex (PPC) plays an important role in multimodal integration (Maravita, Spence, & Driver, 2003). In monkeys, the PPC is a known integration site of somatosensory and visuo-spatial information (Graziano, Cooke, & Taylor, 2000; Sakata, Takaoka, Kawarasaki, & Shibutani, 1973; Hihara et al., 2006; Iwamura, 1998; Iwamura, Iriki, & Tanaka, 1994), with visually responsive neurons in ventral premotor cortex of macaque that can continually update their visual receptive fields to track an effector moving through space (Graziano, Hu, & Gross, 1997). Recent neuroimaging work in humans has implicated similar loci of visual-proprioceptive integration (Limanowski & Blankenburg, 2016). When participants saw a photorealistic virtual arm at a location that was congruent with the position of their real arm, increased activity was observed in posterior parietal cortex, compared to when the virtual and real arm were at incongruent locations. Interestingly, when visual and proprioceptive information were congruent, correlations between integration areas, such as posterior parietal cortex, and primary visual cortex (V1) implied direct communication between these areas (Limanowski & Blankenburg, 2016). A follow-up study investigated hand laterality by looking at congruent and incongruent presentations of both the left and the right hand positioned in their respective hemifields. A cluster in left inferior parietal lobule (IPL) responded preferentially to congruent visuo-proprioceptive hand information for *both* the left and right hand, even with the hands presented in different (the left and right) hemifields (Limanowski & Blankenburg, 2017). These studies highlight cortical sites, like the PPC, at which proprioceptive information coexist with visual representations of both hands.

Importantly, multi-modal interactions have the capacity to alter EVC activity, as has been demonstrated by studies on cross-modal facilitation effects. For example, when tactile stimulation of the hand coincided with a visual stimulus on the same side of space, detection of the visual stimulus was facilitated, and visual cortex activity enhanced (Macaluso et al., 2000). Similarly, several studies have assessed visual cortex excitability by inducing visual phosphenes, which are briefly perceived flashes of light in response to a TMS pulse over EVC. A higher level of cortical excitability is assumed to lower the threshold for eliciting a phosphene (de Graaf, Duecker, Stankevich, Ten Oever, & Sack, 2017). Visual cortex excitability was increased at locations of the visual field that coincided with the location at which a hand was touched (i.e. ‘coincident locations’, Ramos-Estebanez et al., 2007). This effect occurred both with the hands crossed and uncrossed, always showing increased excitability when phosphene location and touch were coincident, irrespective of the hand that was touched (Bolognini & Maravita, 2007). After repetitive 1Hz TMS stimulation of PPC (but not EVC), crossmodal facilitation of EVC still occurred at the coincident location when the hands were uncrossed, but when the hands were crossed it was the noncoincident location that led to increased EVC excitability (i.e. when the tactile stimulus occurred at a location in space corresponding to the ipsilateral hemisphere).

Together, these results suggest that PPC is responsible for binding visual and proprioceptive inputs into a common reference frame that relates to the body, and that top-down influences from parietal multimodal integration areas can directly innervate and impact processing in EVC. In the context of the current experiments, if integrated feedback is send back to EVC, the PPC could indeed provide a reference frame that incorporates information from both hands (represented in both hemispheres).

Interestingly, our data hints at the idea that when processing of a visual stimulus is disrupted with TMS, fading ensues. This trend was particularly apparent in Experiment 2, where TMS tended to make the stationary hands fade. Such TMS induced fading is in line with previous work showing fading after TMS over IPS (Kanai, et al., 2008). Thus, while in one case TMS can have the potential to release cortical inhibition (Ling, Pearson, & Blake, 2009) and thereby increase the perceptual quality of a faded hand, in the other case TMS has the potential to decrease the perceptual quality of a non-faded hand under more standard visual processing conditions (Kanai, et al., 2008). This is in line with the idea that TMS is state dependent, meaning that its impact on behavior results from an interaction with ongoing brain states (Silvanto, Muggleton, & Walsh, 2008).

In our paradigm a sensory conflict arises when a hand is removed from the afterimage, resulting in perceptual fading of the removed hand. We propose that such visual fading results from an attempt to match the visual percept to the proprioceptive experience - via active inhibition at the level of EVC. While this type of fading (through sensory conflict) is likely instantiated via feedback from areas integrating multisensory information (like PPC), continued perceptual fading could be implemented at the level of EVC itself. In this context it is important to note that fading of the afterimage as a whole occurs also in the absence of any sensory conflict, and removing a hand could simply be considered a way to speed up the fading process at a punctate visual location. Future work could address whether fading, and inhibitory processes in EVC, are sustained through continued top-down feedback signals or via local inhibitory processes.

Transients in sensory inputs are generally required for conscious perception, while stable inputs (such as retinally stabilized afterimages) are filtered out of the perceptual experience. Here we demonstrate that TMS over EVC can make a faded image more visible again, signifying a causal role for EVC processes in perceptual fading.

## Acknowledgements

This work was supported by the European Union’s Horizon 2020 research and innovation program under the Marie Sklodowska-Curie Grant Agreement No 743941 to RLR. This work is part of the VICI research program No 453-15-008 awarded to ATS, (partly) financed by the Netherlands Organization for Scientific Research (NWO).

## References

Berlucchi G, Rizzolatti G. (1968). Binocularly driven neurons in visual cortex of split-chiasm cats. Science, 159(3812): 308–10.

Billock, V. A., & Tsou, B. H. (2004). What do catastrophic visual binding failures look like? Trends in Neurosciences, 27(2), 84–89. https://doi.org/10.1016/j.tins.2003.12.003

Blatt, G. J., Andersen, R. A., & Stoner, G. R. (1990). Visual receptive field organization and cortico-cortical connections of the lateral intraparietal area (area LIP) in the macaque. Journal of Comparative Neurology, 299(4), 421–445.

Bolognini, N., & Maravita, A. (2007). Proprioceptive Alignment of Visual and Somatosensory Maps in the Posterior Parietal Cortex. Current Biology, 17(21), 1890–1895. https://doi.org/10.1016/j.cub.2007.09.057

Brainard, D. H. (1997). The Psychophysics Toolbox. Spatial Vision, 10(4), 433–436. https://doi.org/10.1163/156856897X00357

Bross, M. (2000). Emmert’s law in the dark: active and passive proprioceptive effects on positive visual afterimages. Perception, 29(11), 1385–1391.

Carey, D. P., & Allan, K. (1996). A motor signal and visual size perception. Experimental Brain Research, 110(3), 482–486. https://doi.org/10.1007/BF00229148

Carlson, T. A., Alvarez, G., Wu, D., & Verstraten, F. A. J. (2010). Rapid Assimilation of External Objects Into the Body Schema. Psychological Science, 21(7), 1000–1005. https://doi.org/10.1177/0956797610371962

Cavada, C., & Goldman-Rakic, P. S. (1989). Posterior parietal cortex in rhesus monkey: I. Parcellation of areas based on distinctive limbic and sensory corticocortical connections. Journal of Comparative Neurology, 287(4), 393–421.

Darwin, E. (1795). Zoonomia; or the laws of organic life (4th Americ). Philadelphia, PA: Edward Earle.

Davies, P. (1973). Effects of movements upon the appearance and duration of a prolonged visual afterimage: 1. Changes arising from the movement of a portion of the body incorporated in the afterimaged scene. Perception, 2(2), 147–153.

de Graaf, T. A., Duecker, F., Stankevich, Y., Ten Oever, S., & Sack, A. T. (2017). Seeing in the dark: Phosphene thresholds with eyes open versus closed in the absence of visual inputs. Brain Stimulation: Basic, Translational, and Clinical Research in Neuromodulation, 10(4), 828–835.

Ditchburn, R. W., & Ginsborg, B. L. (1952). Vision with a stabilized retinal image. Nature, 170(4314), 36–37.

Graziano, M. S. A., Cooke, D. F., & Taylor, C. S. R. (2000). Coding the Location of the Arm by Sight. Science, 290(5497), 1782–1786. https://doi.org/10.1126/science.290.5497.1782

Graziano, M. S., Hu, X. T., & Gross, C. G. (1997). Visuospatial properties of ventral premotor cortex. Journal of Neurophysiology, 77(5), 2268–2292. https://doi.org/10.1152/jn.1997.77.5.2268

Gregory, R. L., Wallace, J. G., & Campbell, F. W. (1959). Changes in the size and shape of visual after-images observed in complete darkness during changes of position in space. Quarterly Journal of Experimental Psychology, 11(January 2015), 54–55. https://doi.org/10.1080/17470215908416288

Hay, J. C., Pick, H. L., & Ikeda, K. (1965). Visual capture produced by prism spectacles. Psychonomic Science, 2(1-12), 215–216. https://doi.org/10.3758/BF03343413

Hihara, S., Notoya, T., Tanaka, M., Ichinose, S., Ojima, H., Obayashi, S., et al. (2006). Extension of corticocortical afferents into the anterior bank of the intraparietal sulcus by tool-use training in adult monkeys. Neuropsychologia, 44(13), 2636–2646. http://doi.org/10.1016/j.neuropsychologia.2005.11.020

Hogendoorn, H., Kammers, M. P., Carlson, T. A., & Verstraten, F. A. (2009). Being in the dark about your hand: Resolution of visuo-proprioceptive conflict by disowning visible limbs. Neuropsychologia, 47, 2698–2703. https://doi.org/10.1016/j.neuropsychologia.2009.05.014

Ilmoniemi, R. J., Virtanen, J., Ruohonen, J., Karhu, J., Aronen, H. J., Näätänen, R., & Katila, T. (1997).Neuronal responses to magnetic stimulation reveal cortical reactivity and connectivity. NeuroReport, 8(16), 3537–3540. https://doi.org/10.1097/00001756-199711100-00024

Iwamura, Y. (1998). Hierarchical somatosensory processing. Current Opinion in Neurobiology, 8, 522–528.

Iwamura, Y., Iriki, A., & Tanaka, M. (1994). Bilateral hand representation in the postcentral somatosensory cortex. Nature, 369, 554–556.

Kanai, R., Muggleton, N. G., & Walsh, V. (2008). TMS over the intraparietal sulcus induces perceptual fading. Journal of Neurophysiology, 100, 3343–3350. https://doi.org/10.1152/jn.90885.2008

Komatsu, H. (2006). The neural mechanisms of perceptual filling-in. Nature Reviews Neuroscience, 7(3), 220–231.https://doi.org/10.1038/nrn1869

Lewis, J. W., & Van Essen, D. C. (2000). Corticocortical connections of visual, sensorimotor, and multimodal processing areas in the parietal lobe of the macaque monkey. Journal of Comparative Neurology, 428(1), 112–137. https://doi.org/10.1002/1096-9861(20001204)428:1<112::AID-CNE8>3.0.∞;2-9

Limanowski, J., & Blankenburg, F. (2016). Integration of Visual and Proprioceptive Limb Position Information in Human Posterior Parietal, Premotor, and Extrastriate Cortex. Journal of Neuroscience, 36(9), 2582–2589. https://doi.org/10.1523/JNEUROSCI.3987-15.2016

Limanowski, J., & Blankenburg, F. (2017). Posterior parietal cortex evaluates visuoproprioceptive congruence based on brief visual information. Scientific Reports, 7(1), 16659. https://doi.org/10.1038/s41598-017-16848-7

Ling S, Pearson J, Blake R. Dissociation of Neural Mechanisms Underlying Orientation Processing in Humans. Current Biology. 2009; 19(17): 1458–62. https://doi.org/10.1016/j.cub.2009.06.069PMID:19682905

Macaluso, E., Frith, C. D., & Driver, J. (2000). Modulation of Human Visual Cortex by Crossmodal Spatial Attention. Science (New York, NY), 289(5482), 1206–1208. https://doi.org/10.1126/science.289.5482.1206

Maravita, A., Spence, C., & Driver, J. (2003). Multisensory integration and the body schema: Close to hand and within reach. Current Biology, 13(13), 531–539. https://doi.org/10.1016/S0960-9822(03)00449-4

Martinez-Conde, S., Macknik, S. L., & Hubel, D. H. (2004). The role of fixational eye movements in visual perception. Nature Reviews Neuroscience, 5(3), 229–240. https://doi.org/10.1038/nrn1348

Martinez-Conde, S., Otero-Millan, J., & MacKnik, S. L. (2013). The impact of microsaccades on vision: Towards a unified theory of saccadic function. Nature Reviews Neuroscience, 14(2), 83–96. https://doi.org/10.1038/nrn3405

Mendola, J. D., Conner, I. P., Sharma, S., Bahekar, a, & Lemieux, S. (2006). fMRI Measures of perceptual filling-in in the human visual cortex. Journal of Cognitive Neuroscience, 18(3), 363–375. https://doi.org/10.1162/jocn.2006.18.3.363

Meng, M., Remus, D. A., & Tong, F. (2005). Filling-in of visual phantoms in the human brain. Nature neuroscience, 8(9), 1248.

Murray, S. O., Boyaci, H., & Kersten, D. (2006). The representation of perceived angular size in human primary visual cortex. Nature Neuroscience, 9(3), 429–434. https://doi.org/10.1038/nn1641

Rademaker, R. L., Wu, D. A., Bloem, I. M., & Sack, A. T. (2014). Intensive tool-practice and skillfulness facilitate the extension of body representations in humans. Neuropsychologia, 56(1), 196–203. https://doi.org/10.1016/j.neuropsychologia.2014.01.011

Ramos-Estebanez, C., Merabet, L. B., Machii, K., Fregni, F., Thut, G., Wagner, T. A., … Pascual-Leone, A. (2007). Visual Phosphene Perception Modulated by Subthreshold Crossmodal Sensory Stimulation. Journal of Neuroscience, 27(15), 4178–4181. https://doi.org/10.1523/JNEUROSCI.5468-06.2007

Riggs, L. A., Ratliff, F., Cornsweet, J. C., & Cornsweet, T. N. (1953). The disappearance of steadily fixated visual test objects. Journal of the Optical Society of America, 43(6), 495501.

Rossi, S., Hallett, M., Rossini, P. M., & Pascual-Leone, A. (2012). Safety, ethical considerations, and application guidelines for the use of transcranial magnetic stimulation in clinical practice and research. Clinical Neurophysiology, 120(12), 323–330. https://doi.org/10.1016/j.clinph.2009.08.016.Rossi

Sakata, H., Takaoka, Y., Kawarasaki, A., & Shibutani, H. (1973). Somatosensory properties of neurons in the superior parietal cortex (area 5) of the rhesus monkey. Brain Research, 64(C), 85–102. https://doi.org/10.1016/0006-8993(73)90172-8

Sasaki, Y., & Watanabe, T. (2004). The primary visual cortex fills in color. Proceedings of the National Academy of Sciences of the United States of America, 101(52), 18251–18256. https://doi.org/10.1073/pnas.0406293102

Shimojo, S., Kamitani, Y., & Nishida, S. (2001). Afterimage of perceptually filled-in surface. Science, 293, 1677–80. https://doi.org/10.1126/science.1060161

Silvanto J, Muggleton N, Walsh V. (2008). State-dependency in brain stimulation studies of perception and cognition. Trends Cogn Sci., 12(12): 447–54. https://doi.org/10.1016/j.tics.2008.09.004PMID:18951833

Sperandio, I., Chouinard, P. A., & Goodale, M. A. (2012). Retinotopic activity in V1 reflects the perceived and not the retinal size of an afterimage. Nature Neuroscience, 15(4), 540–542. https://doi.org/10.1038/nn.3069

Troxler, D. I. P. V. (1804). Über das Verschwinden gegebener Gegenstände innerhalb unseres Gesichtskreises. Ophthalmologische Bibliothek, 2(2), 1–53.

Wald, G. (1955). The photoreceptor process in vision. American Journal of Ophthalmology, 40(5), 18–41.

